# Dementia Severity Is Associated with Resting-State fMRI Network Responses to Nuisance Regression

**DOI:** 10.64898/2026.11.04.764368

**Authors:** Ildar Gabitov, Aleksandr Volkov, Tamoghna Chattopadhyay, Paul M. Thompson, Yury Maximov

## Abstract

Apparent disease effects in resting-state fMRI may be confounded by the effects of preprocessing, making it essential to understand whether nuisance removal affects patients differently across levels of clinical severity. Yet few studies directly examine whether sensitivity to preprocessing is itself clinically informative. We investigated this in 201 ADNI participants spanning normal cognition, mild cognitive impairment, and dementia. We quantified the coordinated temporal structure of the seven Yeo resting-state networks using a delay-based concentration statistic, comparing estimates after regression of 24 motion parameters and linear trends with estimates obtained after additionally removing five anatomical CompCor components. Adding these components generally reduced the measured temporal concentration, but this reduction was smaller in participants with greater dementia severity (+0.472 percentage points per CDR-SB point). In contrast, the same network measures showed little association with severity before adding CompCor. To determine whether this finding depended on how temporal trends were removed, we repeated the analysis with identical discrete cosine regressors added to both preprocessing designs. The clinical association remained similar (+0.464). Controlled simulations showed that temporal concentration increased with the strength of delayed coupling between networks, even when their individual statistical properties and instantaneous covariance remained unchanged. However, changes in observation amplitude and contamination of nuisance regressors also altered the response to preprocessing without changing the underlying neural signal. These findings suggest that disease-related differences in fMRI may partly reflect differential sensitivity to preprocessing. Systematic sensitivity analyses, including multiverse comparisons across preprocessing strategies, may therefore be essential to distinguish genuine disease effects from preprocessing-dependent associations.

## 1 INTRODUCTION

Reliable clinical interpretation of resting-state fMRI requires us to separate disease-related brain abnormalities from the effects of preprocessing. Motion and physiological noise must be removed, but nuisance regression can also alter meaningful network signals. If these effects covary with disease severity, preprocessing may change the apparent relationship between brain function and clinical symptoms. Understanding this sensitivity is therefore essential for interpreting disease associations.

Prior studies show that nuisance regression can substantially alter measured functional connectivity and its susceptibility to motion artifacts [1, 2, 3]. A systematic evaluation of fMRI pipelines also found that processing choices affect the consistency of functional connectome measures [4] and task-based results [5]. More recently, Kay et al. [6] compared connectivity estimates from lower- and higher-motion portions of each participant’s data to assess motion-related bias in brain–behavior associations. However, it remains unclear if the effects of nuisance regression on the temporal organization of large-scale brain networks vary systematically with clinical severity. This question is important in all developmental, neurological and psychiatric conditions assessed with functional MRI.

We address this question in 201 ADNI participants spanning normal cognition, mild cognitive impairment, and dementia. We measure temporal concentration of activity across the seven Yeo networks and quantify how it changes when five anatomical CompCor components are added to motion and trend regression. We test whether this change is associated with dementia severity, assess its robustness to additional temporal filtering, and use controlled simulations to distinguish sensitivity to delayed network interactions from effects introduced by nuisance signals. This approach treats preprocessing sensitivity as an auxiliary outcome measure, and motivates systematic comparisons across preprocessing choices when interpreting clinical fMRI findings.

## 2 METHODS

### 2.1 Clinical Cohort Selection

We analyzed resting-state fMRI from the Alzheimer’s Disease Neuroimaging Initiative (ADNI) [7], comprising 4,347 BOLD series from 4,257 visits in 1,764 participants. Eligible visits required linked clinical and imaging records, a diagnosis recorded within 90 days, and a corresponding anatomical MRI scan. To reduce differences in sampling and acquisition across diagnostic groups, we matched all 90 eligible dementia histories to 90 cognitively normal (CN) and 90 mild cognitive impairment (MCI) histories. Matching accounted for demographic and acquisition characteristics and required identical numbers of eligible visits, yielding 270 participants and 732 visits.

We further required a Clinical Dementia Rating Sum of Boxes (CDR-SB) assessment within 90 days of imaging. Retaining the first of 615 clinically eligible visits per participant left 267 individuals. Because our temporal analysis uses fixed sample delays, we restricted scans to a 3-second repetition time (TR) to ensure comparable temporal scales. The final cohort comprised 201 individuals: 64 CN, 70 MCI, and 67 with dementia. We analyzed CDR-SB as a continuous severity measure [8] to test whether nuisance-regression sensitivity varies with symptom burden beyond diagnostic category.

### 2.2 Nuisance regression and network dynamics

To measure how preprocessing alters network dynamics, we analyzed each scan using two nuisance-regression models. Let *X* ∈*R*^*N*×400^ contain *N* retained samples of the Schaefer parcel time series [9] and *W* ∈*R*^400×7^ average them into the seven Yeo networks. The baseline model, *D*_0_, removes an intercept, linear trend, and 24 motion regressors. The augmented model, [*D*_0_, *A*], additionally removes five anatomical CompCor components, collected in *A* and selected by their leading singular values from the combined anatomical mask [10]. Both models regress all nuisance terms simultaneously, avoiding interactions introduced by sequential filtering [11]. For an orthonormal basis *U*_*D*_ of either nuisance design, we have:

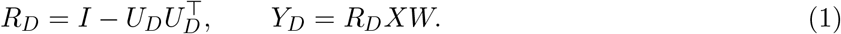

Here, *I* is the *N* × *N* identity, *R*_*D*_ is the residual projector, and *Y*_*D*_ contains the resulting network time series. Both designs use joint projection. Each resulting network time series is centered and normalized to unit root-mean-square amplitude, so the analysis emphasizes temporal organization rather than signal magnitude. To capture interactions across time, we construct a delay matrix *H*_*L*_ by stacking *L* = 10 consecutive samples from each network, producing a 70 × (*N* −9) matrix spanning 27 seconds [12]. We measure the fraction of its total variance explained by the two leading singular modes. To distinguish coordinated temporal structure from the individual networks’ frequency spectra, we subtract the average concentration from 199 phase-randomized reference series [13]:

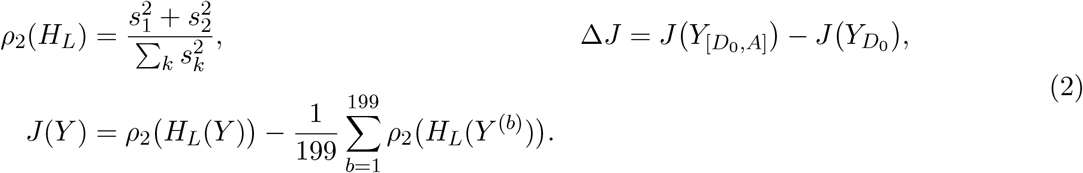

Here, *s*_*k*_ are the singular values of *H*_*L*_ in descending order, with the sum over all k; *Y* denotes either normalized network-series matrix and *Y* ^*b*^ its *b*-th phase-randomized reference. The reference series preserve each network’s Fourier-amplitude spectrum while independently randomizing its phases. Our primary outcome, Δ*J*, measures how much additional anatomical nuisance removal changes temporal concentration. We report 100*J* and 100Δ*J* in percentage points (pp). To test whether this response depends on the treatment of slow temporal fluctuations, we repeat both regressions after adding identical discrete cosine transform (DCT) regressors. These are fitted jointly with the other nuisance terms, while keeping the five anatomical components unchanged.

### 2.3 Clinical Associations and Statistical Inference

We tested whether the change in temporal concentration, Δ*J*, was associated with dementia severity measured by CDR-SB. Models adjusted for age, sex, scan length, acquisition characteristics, site, scanner, motion, and signal-quality measures, including an age-proxy indicator. Using the common covariate design *Z*, severity vector *c* and residualized severity *z* = *R*_*Z*_*c*, where *R*_*Z*_ projects off the column space of *Z*, the paired coefficient is

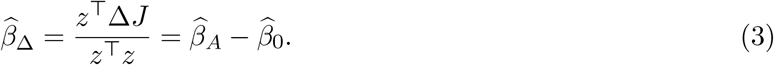

Here, Δ*J* collects 201 participant responses;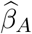, 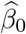 denote adjusted severity slopes for augmented and baseline outcomes, respectively. This tests whether preprocessing sensitivity varies with severity while accounting for within-scan correlations. We estimate uncertainty using HC3 standard errors [14] and a restricted bootstrap (20,000 draws; HC3 studentization [15]). After covariate adjustment, 14 observations do not inform the severity coefficient.

As the cohort contributed to earlier exploratory analyses, we regard any clinical association as a discovery rather than an independent confirmation. The additional DCT analysis tests its sensitivity to temporal preprocessing. We also evaluate bootstrap calibration under simulated constant and unequal error variances to assess the reliability of the nominal significance levels.

## 3 RESULTS

### 3.1 Preprocessing Sensitivity Varies with Dementia Severity

Adding anatomical CompCor components generally reduced temporal concentration across the seven Yeo networks, but less so with greater dementia severity. After adjusting for demographic, acquisition, and motion covariates, each additional CDR-SB point was associated with a 0.472 percentage-point increase in paired response (HC3 SE = 0.247, partial r = 0.233; nominal bootstrap p = 0.0148). The association remained similar after adding identical temporal DCT regressors to both preprocessing models (slope = 0.464, HC3 SE = 0.221, partial r = 0.237; nominal p = 0.0076; Table 1, Fig. 1), indicating consistency across temporal preprocessing.

**Table 1.** Association of dementia severity (CDR-SB) with temporal concentration changes after additional anatomical nuisance regression (201 participants). Slopes and HC3 SEs are in percentage points per CDR-SB point; bootstrap p-values are nominal and approximate.

| Design | Slope | HC3 SE | Partial r | Nominal p |
|---|---|---|---|---|
| Original | 0.4722 | 0.2470 | 0.2332 | 0.0148 |
| Common DCT | 0.4641 | 0.2210 | 0.2365 | 0.0076 |

**Figure 1.**
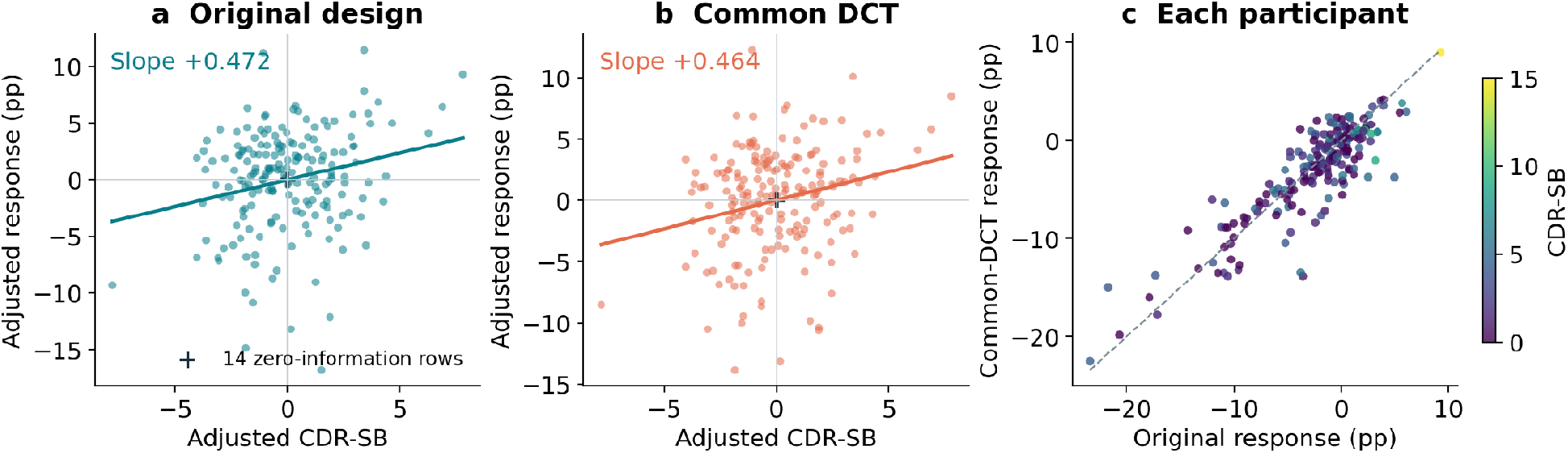
Association between dementia severity and sensitivity to nuisance regression in 201 ADNI participants. (a,b) Changes in temporal concentration after adding anatomical CompCor, plotted against CDR-SB, before (a) and after (b) additional temporal DCT regression. Both variables are adjusted for demographic, acquisition, and motion-related covariates; lines show the fitted associations. Fourteen observations with no remaining information about the severity coefficient overlap at the origin. (c) Comparison of individual preprocessing responses with and without DCT regression, colored by dementia severity (CDR-SB).

The clinical association reflected sensitivity to nuisance regression rather than temporal concentration itself. Before CompCor, concentration was nearly unrelated to CDR-SB (slope = −0.0443 pp, partial r = −0.019, nominal p = 0.816). CompCor reduced concentration by 2.6 pp on average, with smaller reductions at greater dementia severity. This pattern persisted with additional temporal DCT regression, which increased the average reduction to 3.1 pp. Thus, the association reflected responses to nuisance removal rather than baseline concentration.

Because our measure subtracts phase-randomized reference concentration, we tested whether this correction explained the association. Both observed and reference concentrations varied with severity, but their differences retained positive clinical slopes of 0.47 and 0.46 pp across the two preprocessing designs. Reference-generation uncertainty was negligible relative to between-participant variability (Monte Carlo SE = 0.0038 pp with common DCT).

### 3.2 Reliability of Bootstrap Significance Estimates

Given the exploratory clinical analysis, we assessed bootstrap significance calibration using 1,000 simulated datasets under constant error variance or variance varying with scan length. With 20,000 bootstrap draws, false-positive rates at the nominal 1% threshold were 0.6% and 1.9%, respectively. The latter exceeded the expected 1% (95% CI: 1.15–2.95%), indicating overstated significance under unequal variance. Rejection rates at 5% and 10% remained consistent with nominal levels. These findings do not affect clinical association estimates or their stability across preprocessing designs but warrant caution when interpreting small nominal bootstrap p-values near 1%.

### 3.3 Controlled Simulations of Network Dynamics and Preprocessing Effects

To interpret the clinical association, we used controlled simulations to determine what the temporal concentration measure detects and whether changes in nuisance signals could produce similar effects without changes in the underlying brain dynamics.

### 3.3.1 Detecting Delayed Network Interactions

We first tested whether temporal concentration detects interactions between networks that would be missed by conventional zero-lag connectivity. We generated seven Gaussian time series, introducing delayed coupling between two while keeping each network’s individual distribution and temporal spectrum unchanged:

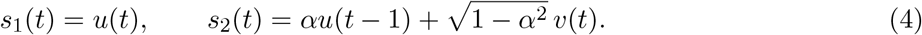

Here, *t* indexes time samples, *s*_1_ and *s*_2_ denote two simulated network signals, *u* and *v* are independent standard Gaussian white noise, and *α* ∈[0, 1] controls coupling strength. The population covariance at zero lag remains the identity matrix, regardless of *α*, but the delay representation captures the interaction. Its two leading eigenvalues increase to 1 + *α*, giving a theoretical leading-two concentration of 2(1 + *α*)*/*70 (seven networks, 10-time-point delay).

We tested coupling strengths of 0, 0.5, and 1 across three scan lengths (97, 197, and 788 frames), with 64 independent realizations per length. Increasing coupling from zero to one raised excess temporal concentration by 1.3, 1.7, and 2.2 percentage points, respectively (Fig. 2). The effect persisted when circular delays eliminated boundary effects, confirming that it reflects delayed interactions rather than the construction of the delay matrix. Thus, temporal concentration can detect network relationships that are absent from the population’s instantaneous covariance.

**Figure 2.**
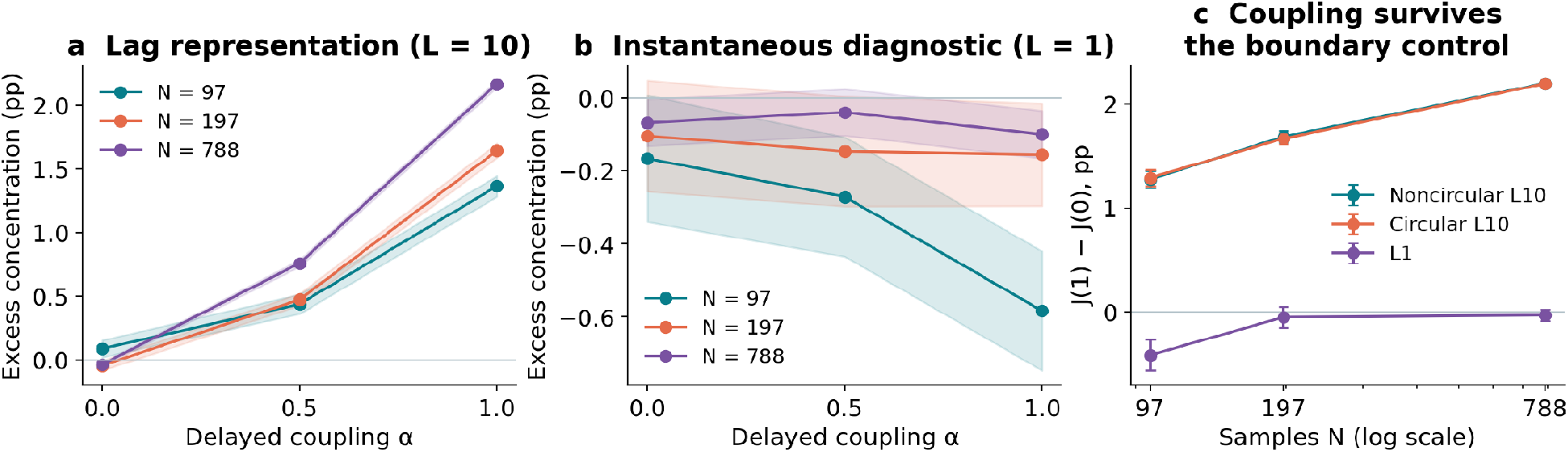
Detecting delayed interactions between simulated brain networks. (a) Temporal concentration increases with delayed coupling despite unchanged individual network statistics and population zero-lag correlations. Results cover three time-series lengths. (b) The simultaneous-observation measure varies with sample length despite unchanged population covariance. (c) The increase from zero to full coupling, J(1) − J(0), persists with circular delays that eliminate boundary effects, ruling out delay-matrix construction artifacts. Shaded bands indicate one standard error across 64 simulations. L10 and L1 denote L = 10 and L = 1, respectively.

#### 3.3.2 Effects of Observation and Nuisance Regression

We next asked whether preprocessing sensitivity could change with the underlying neural signal fixed. Simulations varied observation amplitude, nuisance-proxy quality, shared latent activity, and nuisance-regressor contamination, using two scan lengths and 16 independent realizations per condition (Fig. 3).

**Figure 3.**
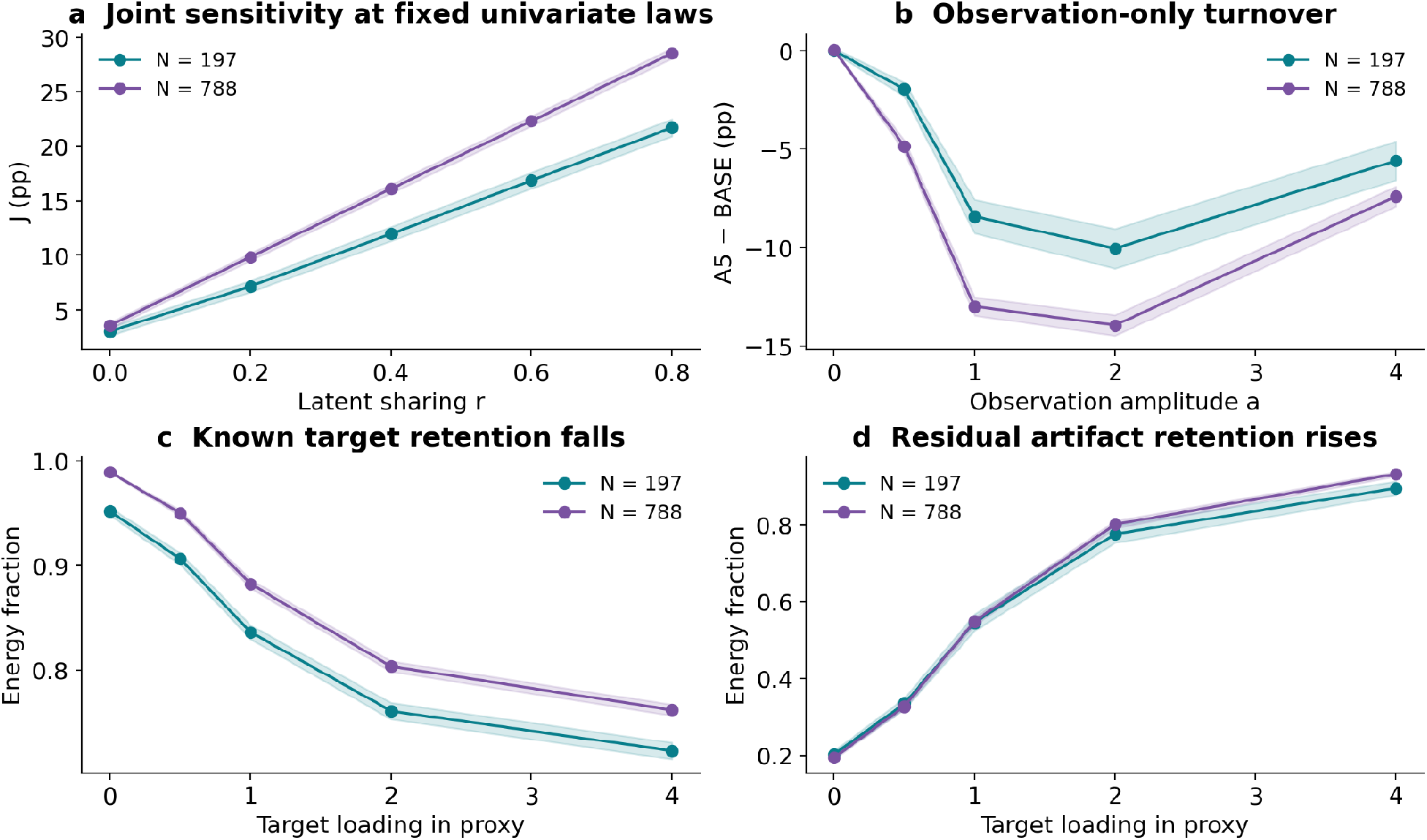
How neural signals and nuisance regression affect temporal concentration. (a) Greater shared activity between networks raises temporal concentration without changing individual network statistics. (b) Changing nuisance-signal amplitude alters preprocessing responses despite unchanged underlying neural activity. (c,d) Increasing target-signal contamination of nuisance regressors inadvertently removes more neural signal (c) while leaving more artifacts in processed data (d). Thus, nuisance regression can remove meaningful brain signals while failing to eliminate artifacts, potentially distorting clinical associations. Results are shown for two time-series lengths; shaded bands indicate one standard error across 16 simulations.

Increasing shared latent activity raised temporal concentration from 3.041 to 21.711 pp at 197 frames, despite unchanged individual-channel distributions. However, changing observation amplitude alone also altered the paired preprocessing response, which varied nonmonotonically: increasing amplitude from two to four raised the response by 4.460 pp at 197 frames and 6.543 pp at 788 frames. When nuisance regressors contained increasing amounts of the target signal, target-energy retention fell from 95.2% to 72.3%, while artifact retention rose from 20.2% to 89.6%.

These experiments establish two important properties. Temporal concentration can detect delayed network organization that conventional connectivity measures may miss, but its response to nuisance regression also depends on observation conditions and how well nuisance signals are separated from the target. Consequently, the clinical association cannot be attributed uniquely to altered neural dynamics without further analysis. The *L* = 1 diagnostic varies despite fixed population covariance. A population constraint does not fix the distribution of a finite estimate.

## 4 DISCUSSION & CONCLUSION

Reliable interpretation of clinical fMRI requires understanding how brain measurements change when nuisance signals are removed. We found that anatomical nuisance regression reduced temporal concentration less in participants with greater dementia severity, even though the original measure showed little clinical association. This relationship persisted after additional temporal filtering, suggesting that preprocessing sensitivity is crucial to understand in clinical studies that use fMRI. Our simulations showed that temporal concentration can detect interactions between brain networks that occur with a time delay and are missed by conventional connectivity measures. However, changes in nuisance signals and the effectiveness of their removal also altered the results, even when the underlying neural activity was unchanged. Thus, clinical associations may reflect genuine differences in brain function, and differences in how preprocessing affects the data.

These exploratory findings require independent validation, particularly given the modest sample size and limitations of bootstrap calibration. Future work should identify the neural and physiological sources of preprocessing sensitivity and test clinical associations across a wider range of processing strategies. Multiverse analyses could establish which findings remain robust to these choices. More broadly, preprocessing sensitivity should be measured and reported, to distinguish genuine disease-related brain alterations from effects introduced or modified by preprocessing.

## 5 CODE AND DATA AVAILABILITY

Code and data for the paper are available at https://github.com/yury-rd/clinical-response

## 6 COMPLIANCE WITH ETHICAL STANDARDS

This secondary analysis used existing public ADNI data^1^, which was collected with IRB approval for the initial study. Restricted data were not redistributed.

## Acknowledgments

Data used in preparation of this article were obtained from the Alzheimer’s Disease Neuroimaging Initiative (ADNI) database (adni.loni.usc.edu). As such, the investigators within the ADNI contributed to the design and implementation of ADNI and/or provided data but did not participate in the analysis or writing of this report. A complete listing of ADNI investigators can be found in the ADNI investigator acknowledgment list.

The ADNI imaging and clinical data came from the Alzheimer’s Disease Neuroimaging Initiative database. ADNI began in 2003 as a public-private partnership led by Michael W. Weiner, MD. It brings together longitudinal imaging, biomarkers, and clinical and neuropsychological assessments to study progression from mild cognitive impairment to Alzheimer’s disease.

The National Institute on Aging funds ADNI data collection and sharing through National Institutes of Health grant U19 AG024904 to the Northern California Institute for Research and Education. ADNI has also received support from the National Institute of Biomedical Imaging and Bioengineering, the Canadian Institutes of Health Research, and private contributors through the Foundation for the National Institutes of Health. These contributors include AbbVie, Alzheimer’s Association, Alzheimer’s Drug Discovery Foundation, Araclon Biotech, BioClinica, Inc., Biogen, Bristol-Myers Squibb Company, CereSpir, Inc., Cogstate, Eisai Inc., Elan Pharmaceuticals, Inc., Eli Lilly and Company, EuroImmun, F. Hoffmann-La Roche Ltd and its affiliated company Genentech, Inc., Fujirebio, GE Healthcare, IXICO Ltd., Janssen Alzheimer Immunotherapy Research & Development, LLC., Johnson & Johnson Pharmaceutical Research & Development LLC., Lumosity, Lundbeck, Merck & Co., Inc., Meso Scale Diagnostics, LLC., NeuroRx Research, Neurotrack Technologies, Novartis Pharmaceuticals Corporation, Pfizer Inc., Piramal Imaging, Servier, Takeda Pharmaceutical Company, and Transition Therapeutics.

## Footnotes

1 https://adni.loni.usc.edu/data-samples/adni-data/

